# An Entropy Approach for Choosing Gene Expression Cutoff

**DOI:** 10.1101/2022.05.05.490711

**Authors:** Hy Vuong, Tung Nguyen, Huy Nguyen, Thao Truong, Son Pham

## Abstract

Annotating cell types using single-cell transcriptome data usually requires binarizing the expression data to distinguish between the background noise vs. real expression or low expression vs. high expression cases. A common approach is choosing a “reasonable” cutoff value, but it remains unclear how to choose it. In this work, we describe a simple yet effective approach for finding this threshold value.

## 1 Introduction

A common procedure to annotate cell types in a single-cell RNA-seq study is to first perform graph-based clustering, and further check the expression of some marker genes in each cluster. In some cases, scientists need to distinguish between the real expression of a gene vs the background expression. In other cases, they need to know if a gene expresses highly in one cluster and lowly in another cluster (e.g., NK *CD*56^*bright*^ and NK *CD*56^*dim*^). This requires choosing a threshold to binarize the expression data. It remains unclear how to choose this threshold. Here, we propose to binarize the data in a way that minimizes the clustering information loss. Below, we describe the formulation in detail.

## 2 Method

*H* is the entropy [2] (information theory) function

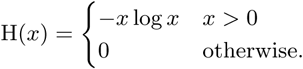

Denote *C* as the set of cells. Each cell *c ∈ C* has an expression value denoted by *η*(*c*).

Given a cut-off threshold *t*, we denote the set of expressed cells *E*_*t*_, and set of non-expressed cells *Ē*_*t*_ as

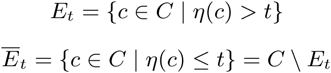

We also denote a partition (clustering) of cells as a set *P* of *k* disjoint cell subsets,

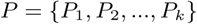

Where, 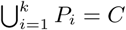, and ∀*i*≠ *j, P*_*i*_ ∩ *P*_*j*_ = ∅

Finally, we define our optimal cutoff threshold as

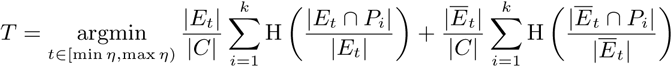

The threshold *t* should split C into two non-empty sets *E*_*t*_ and *Ē*_*t*_. Hence, min *η ≤ t <* max *η*. Intuitively, this value of *t* minimizes the clustering information loss.

## 3 Results

We applied this thresholding method to binarize the expression data of gene CD8A from GSE108989 [3]. This data contains 11,138 single T cells from 12 patients with colorectal cancer with RNA sequencing and TCR tracking (STARTRAC) indices. The annotation result is highly consistent with authors’ annotations for CD8 subtypes (See Fig.1). Fig. 2 shows the violin plots for the expression of each cluster, before and after cutoff. The optimal cutoff value is chosen at the point where the entropy function reaches its minimum (Fig. 3).

**Figure 1:**
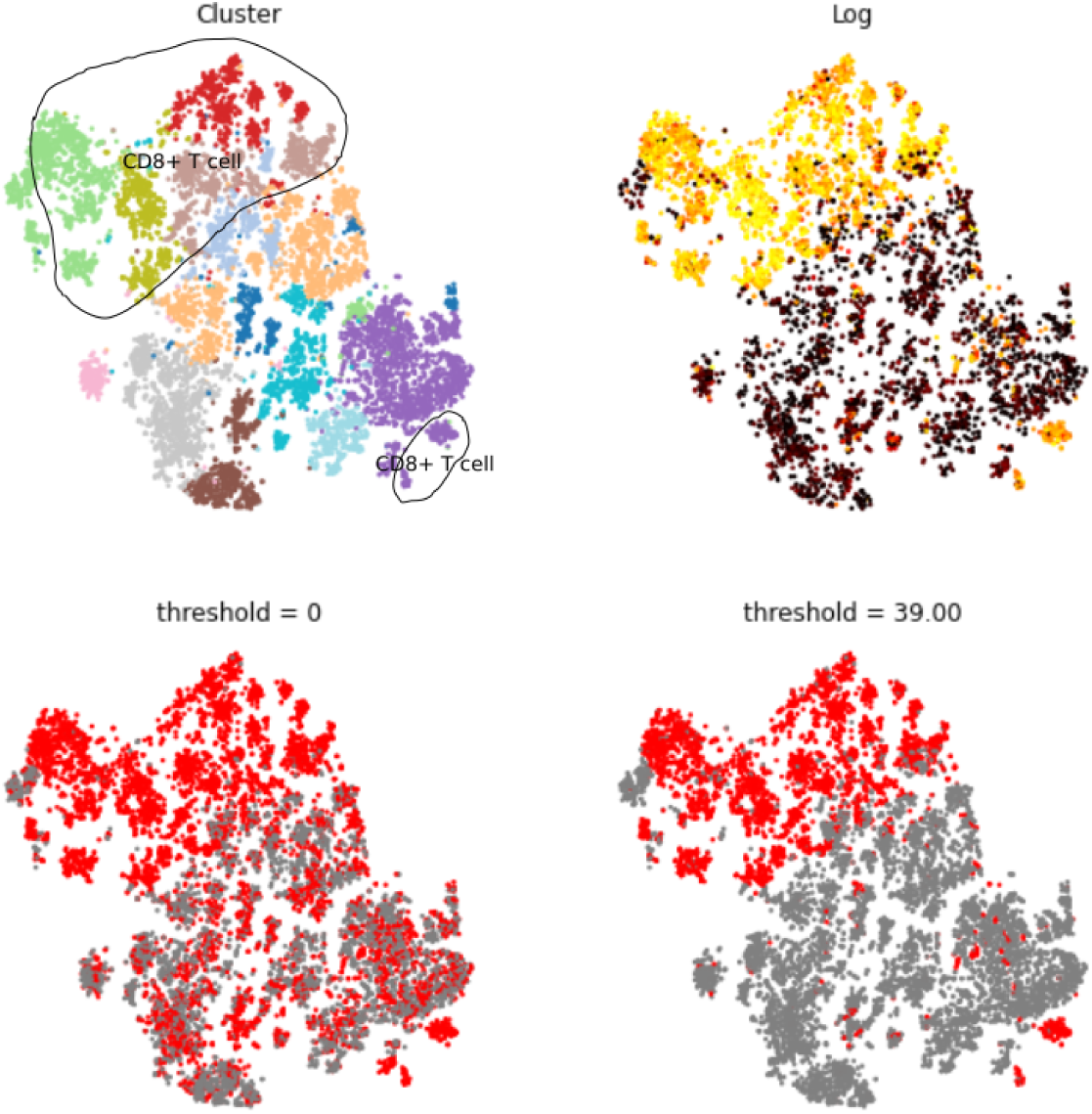
t-SNE plot for 11,138 cells. (a) The plot is colored by graph-based clustering with CD8 subtypes in circles. (b) The plot is colored by the log of the raw expression values. (c) The plot shows the expression of CD8A where red dots correspond to cells with CD8A expression *>* 0, and gray dots correspond to cells with zero CD8A expression.(d) The plot is colored by using binarized version of CD8A gene, using the optimal cutoff value derived from Fig. 3

**Figure 2:**
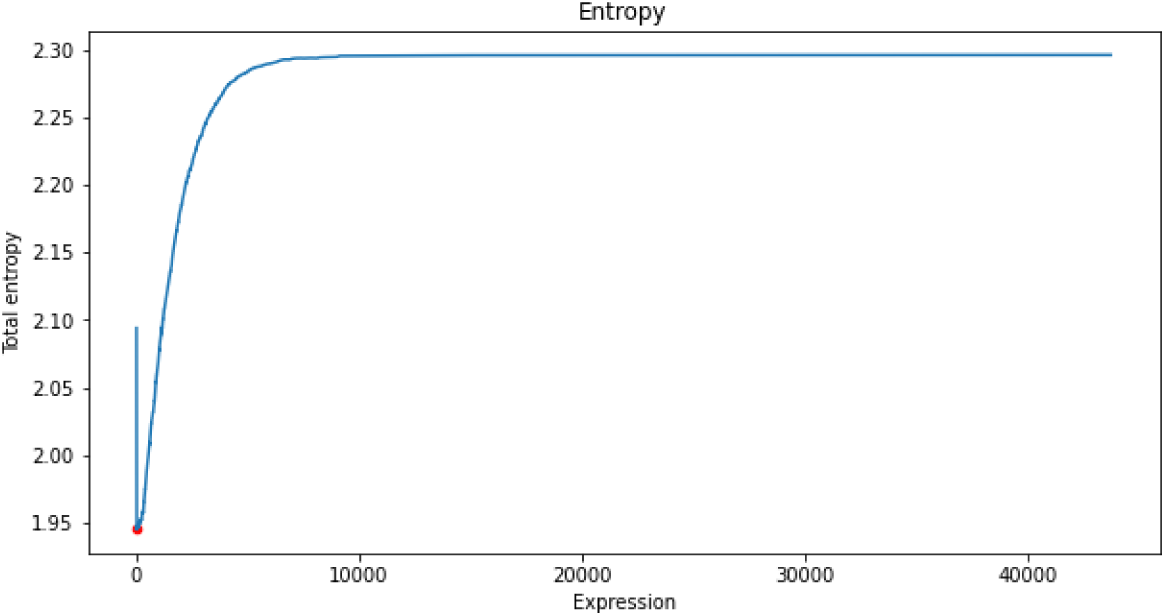
Entropy plot for CD8A in dataset GSE108989. The best threshold to minimize total entropy is T = 39.

**Figure 3:**
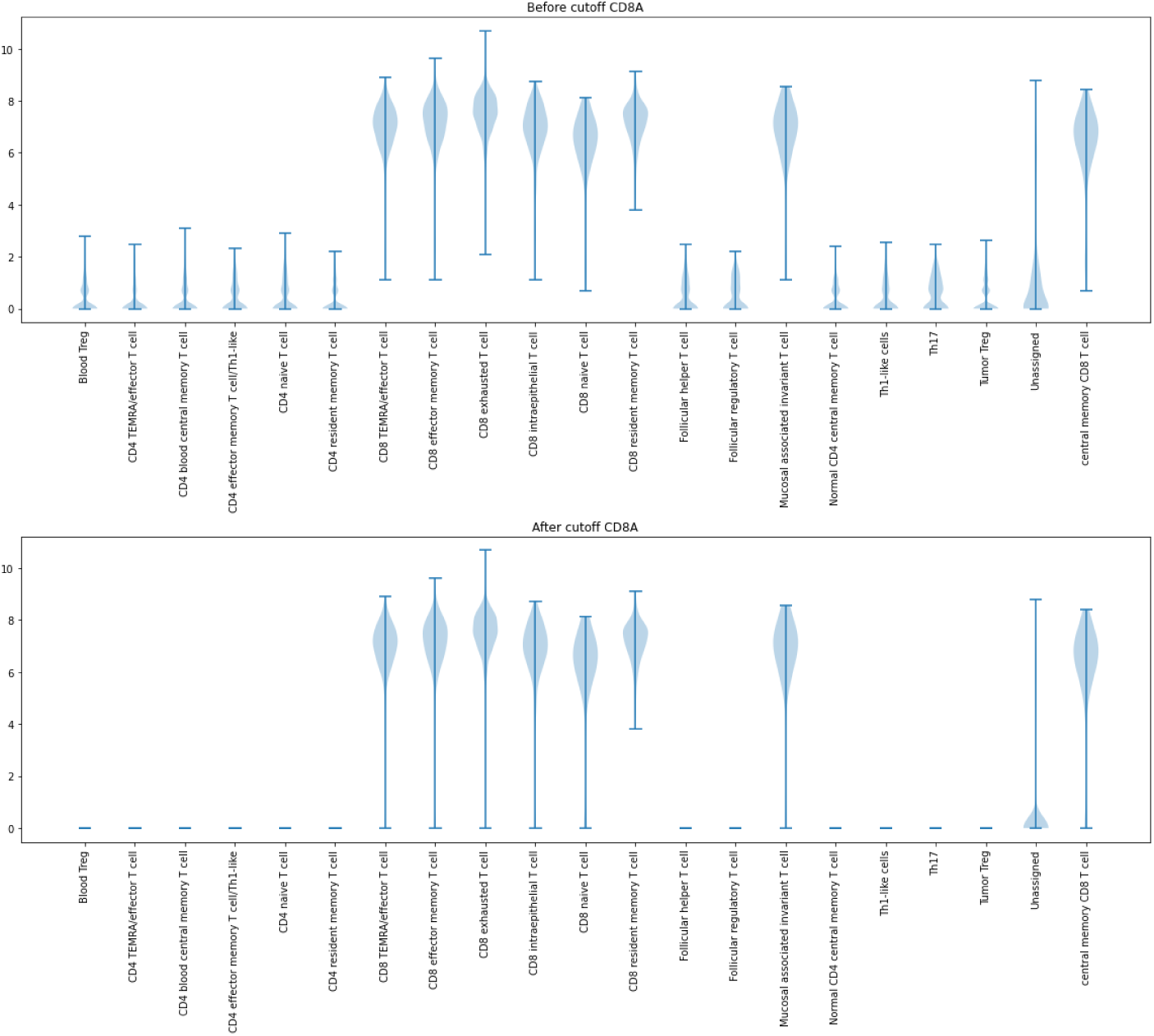
Violin plot for CD8A expression of 11,138 cells from dataset GSE108989 split by author’s annotation (cell types). (a) The violin plot of log raw expression values. (b) The violin plot of log raw expression values using cutoff with threshold T = 39 (raw value).

## 4 Discussion

We presented a simple approach for selecting a threshold to binarize single-cell RNA-seq expression data that minimizes the clustering information loss. This idea is also similar to how a decision tree [1] calculates the entropy of each feature after every split. This approach (when combined with kNN smoothing) leads to consistent annotation results with manual cell type annotations from many published papers and can be helpful for future cell type annotation tasks.

